# Mammalian olfaction is a high temporal bandwidth sense

**DOI:** 10.1101/570689

**Authors:** Andrew Erskine, Tobias Ackels, Debanjan Dasgupta, Izumi Fukunaga, Andreas T. Schaefer

## Abstract

Odours are transported in turbulent plumes resulting locally in highly fluctuating odour concentration (Celani et al., 2014; Murlis et al., 1992; Mylne and Mason, 1991; Shraiman and Siggia, 2000). Yet, whether mammals can make use of the ensuing temporal structure (Celani et al., 2014; Crimaldi and Koseff, 2001; Murlis et al., 1992; Mylne and Mason, 1991; Schmuker et al., 2016; Vickers, 2000) to extract information about the olfactory environment remains unknown. Here, we use dual-energy photoionisation recording with >300 Hz bandwidth to simultaneously determine odour concentrations of two odours in air. We show that temporal correlation of odour concentrations reliably predicts whether odorants emerge from the same or different sources in normal turbulent environments outside and in laboratory conditions. To replicate natural odour dynamics in a reproducible manner we developed a multichannel odour delivery device allowing presentation of several odours with 10 ms temporal resolution. Integrating this device in an automated operant conditioning system we demonstrate that mice can reliably discriminate the correlation structure of odours at frequencies of up to 40 Hz. Consistent with this finding, output neurons in the olfactory bulb show segregated responses depending on the correlation of odour stimuli with populations of 10s of neurons sufficient to reach behavioural performance. Our work thus demonstrates that mammals can perceive temporal structure in odour stimuli at surprisingly fast timescales. This in turn might be useful for key behavioural challenges (Jacobs, 2012) such as odour source separation (Hopfield, 1991), figure-ground separation (Rokni et al., 2014) or odour localisation (Vergassola et al., 2007; Vickers, 2000).

---

The turbulent nature of air as well as water flow results in complex temporal fluctuations of odour concentrations that depend on the distance and direction of odour sources (Atema, 1995; Baker et al., 1998; Balkovsky and Shraiman, 2002; Celani et al., 2014; Crimaldi and Koseff, 2001; Fackrell and Robins, 1982; Justus et al., 2002; Mafraneto and Carde, 1994; Moore and Atema, 1991; Murlis et al., 1992; Murlis et al., 2000; Mylne and Mason, 1991; Schmuker et al., 2016; Vergassola et al., 2007; Vickers, 2000; Vickers et al., 2001; Weissburg et al., 2012; Weissburg et al., 2002). Natural odours consist of multiple different types of molecules (Mori and Yoshihara, 1995), and a typical olfactory scene contains several of such sources (Rokni et al., 2014), with possibly overlapping chemical compositions. Thus, in order to identify naturally complex odours, the first challenge the olfactory system faces is to separate odour sources and possibly attribute different chemicals to different sources (Hopfield, 1991; Jacobs, 2012). Motivated by the turbulent nature of odour transport, it was suggested that the temporal structure of odour concentration fluctuations might contain information regarding odour source (Hopfield, 1991) - i.e. that chemicals belonging to the same sources would co-fluctuate in concentration, whilst chemicals belonging to different sources would be largely uncorrelated. Therefore if animals can detect these correlation structures they would easily be able to perceive which odours arise from the same object.

In order to test this in air, we developed a dual-energy fast photoionisation detection (defPID) approach to measure odour concentrations of two odours simultaneously with high temporal bandwidth (methods, **Fig 1a-c, Supp Figs 1.1–1.2**). When an odour was presented in a laboratory environment with artificially generated turbulence (**Fig 1a**), odour concentration was indeed highly fluctuating with a spectrum extending beyond 40 Hz (**Supp Fig 1.1f**). If two odours were presented from the same source, these fluctuations were highly correlated (**Fig 1a,b**). When we separated odour sources and presented the two odours from as little as 50 cm apart, however, odour dynamics were almost completely uncorrelated (**Fig 1a,b**) with intermediate correlations for closer distances (**Fig 1b**). This pattern of almost perfect correlation for same source and virtually uncorrelated dynamics for sources separated by as little as 50 cm was maintained at closer and further distances between odour source and sensor (**Supp Fig 1.2a,b**), independent of the odours used (**Supp Fig 1.2c**) and mirrored in a natural environment (outside, **Fig 1c**). Thus, the correlation structure of odorant concentration fluctuations indeed contains reliable information about the distribution of odour sources in air. Can mice make use of this information? In order to reproducibly probe the behavioural response to fluctuating stimuli we constructed a high bandwidth odour delivery device based on high-speed solenoid valves operating at >1 kHz (methods, **Fig 1d-h, Supp Fig 1.3–1.4**). Odour pulses could be generated at frequencies of 40 Hz with minimal loss in fidelity (**Fig 1e,f**) for different odours (**Supp Fig 1.3–1.4**). Pulse-width modulation at sub-millisecond time scales makes rapid and reproducible concentration adjustment possible (**Fig 1.3c**) allowing us to reliably emulate the dynamic features of natural plumes. A multiple channel version of the odour delivery device in turn could reliably produce odour pulses with different temporal correlation structures (**Fig 1h**).

**Figure 1:**
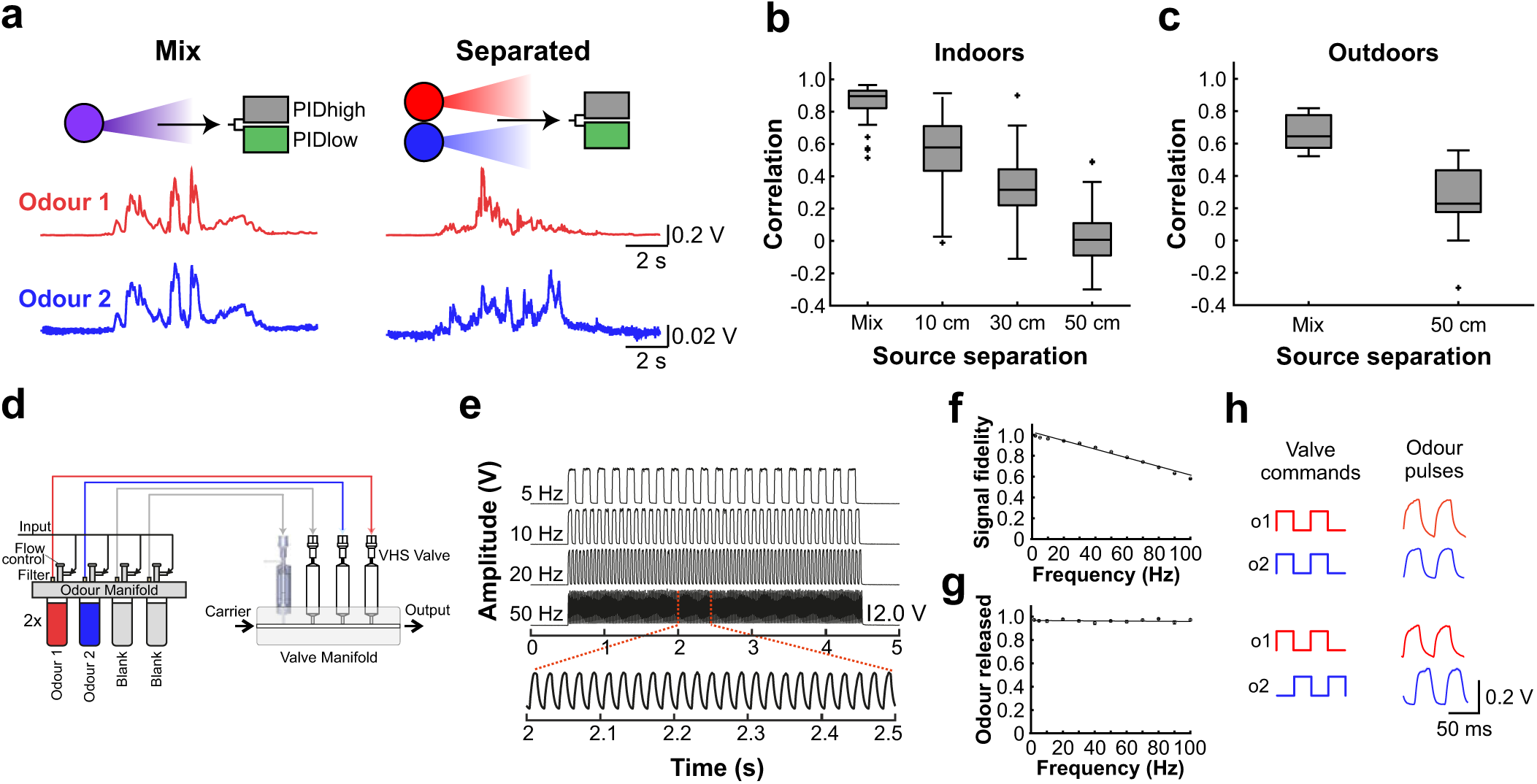
Dual-energy fast photoionisation detection (defPID) and high bandwidth odour delivery. **a,** Simultaneous measurement of two odours (Odour 1: α-Terpinene, Odour 2: Ethyl butyrate) presented either from the same source or separated from each other by 50 cm using two PIDs with bulbs that have different ionisation energies. **b,** Average correlation (mean +/− SD) over all recordings (n = 61 for Mix, n = 71 for each individual distance) for mixed sources and sources separated by multiple distances in a controlled laboratory environment. **c,** Average correlation (mean +/− SD) for mixed sources and sources separated by 50 cm measured outdoors (n = 7, 10). **d,** Schematic of the multi-channel high bandwidth odour delivery device. **e,** Representative odour pulse recordings at command frequencies between 5 and 50 Hz. **f,** Relationship of frequency and odour pulse signal fidelity (for calculation see methods section) and **g,** total amount of odour released (n = 5 repeats for each condition, mean +/− SEM). **h,** Exemplary traces of decomposed correlated and anti-correlated odour pulses presented at 20 Hz.

In order to assess whether mice could discriminate between such stimuli at different frequencies, we developed an automatic operant conditioning system (“AutonoMouse”, **Fig 2a, Supp Video 1** (Erskine et al., 2019)), where cohorts of up to 25 mice could be trained in operant conditioning tasks simultaneously. Mice in the system were group-housed and RFID-tagged for individual identification, remaining in the apparatus for periods of up to 18 months. Food was freely accessible but water could be obtained only by completion of go/no-go operant conditioning tasks. Mice indeed readily learned to differentially respond to correlated or anti-correlated odours (**Fig 2b-d**). Gradually increasing the correlation frequency showed that animals were capable of reliably detecting the correlation structure of the stimuli at frequencies of up to 40 Hz (**Fig 2d-f**). As a population, animal performance decreased by approximately 5 % per octave with performance significantly above chance at frequencies of up to 40 Hz (n=33 mice in two cohorts of 14 and 19 mice, **Fig 2f, Supp Fig 2.3**). To mitigate the risk of animals using non-intended cues for discrimination, odours were presented from changing combinations of valves for each trial (**Fig 2c, Supp Fig 2.1**), odour flow was carefully calibrated and additionally varied randomly between trials such that neither flow nor average concentration provided any information about the nature of the stimulus (**Supp Fig 2.1e,f**). Consistent with this, when valve identities were scrambled, animals performed at chance (orange, **Fig 2f, Supp Fig 2.3**). Finally, when odour presentation was changed to a new set of valves (Abraham et al., 2004), performance levels were maintained (**Supp Fig 2.1d,** inset **Fig 2d**), indicating that only intended cues (the temporal structure of odours) were used for discrimination.

**Figure 2:**
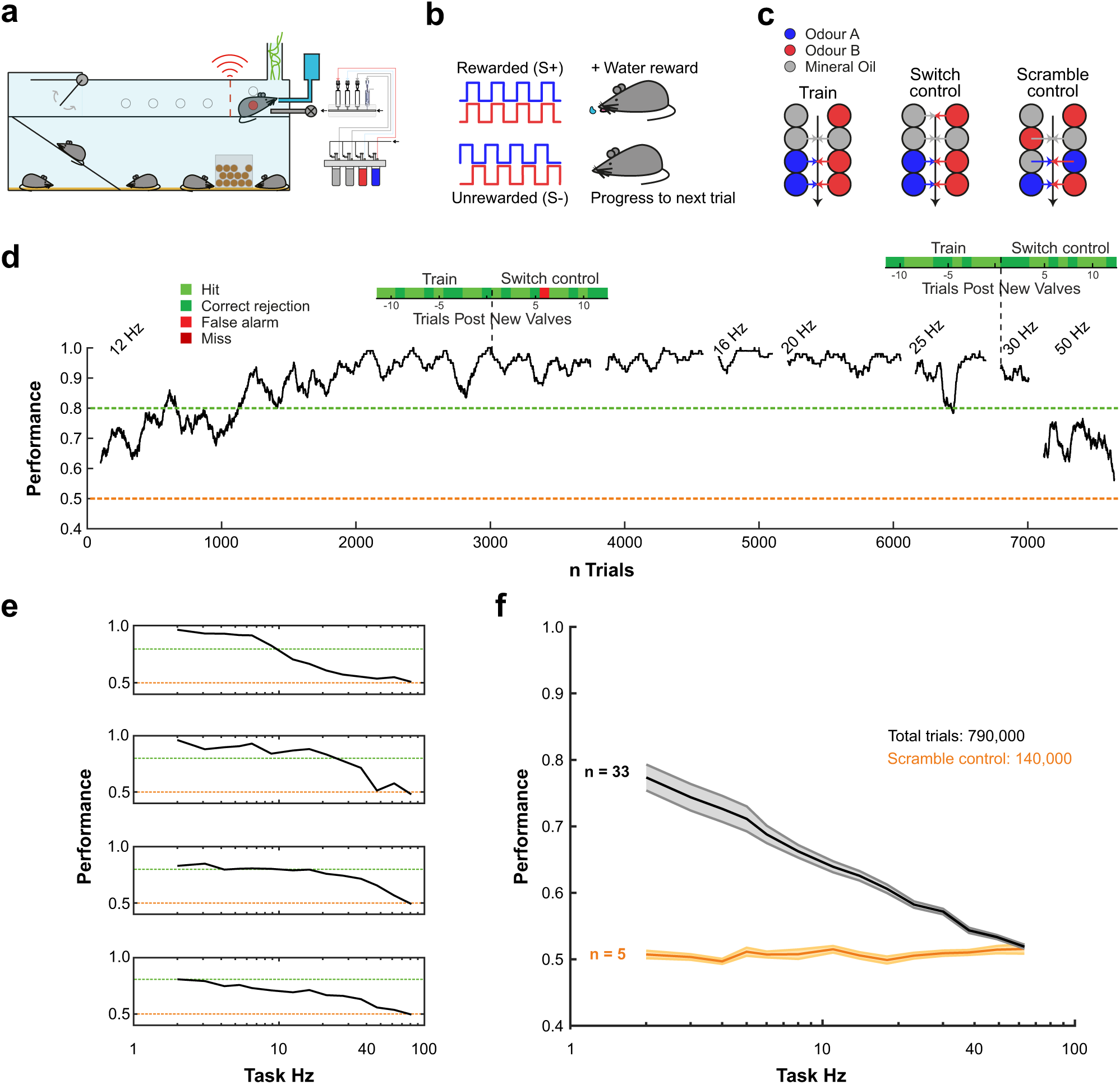
Animals learn to discriminate odour correlation structure. **a,** Schematic of the automated operant conditioning system (“AutonoMouse”) housing cohorts of up to 25 animals. **b,** Schematic of the discrimination stimuli; mice were trained to discriminate between two odours presented simultaneously in either a correlated (top) or anti-correlated (bottom) fashion. Choice was indicated by licking behaviour in a standard go/no-go paradigm. **c,** Schematic of valve combinations for stimulus production. Train: a subset of 6 valves is used to produce the stimulus through varying valve combinations. Switch control: two extra valves are introduced and odour presentation switched over to the newly introduced valves. Scramble control: valve maps (represented by arrow colour) are maintained compared to the train condition but odour vial positions are scrambled resulting in odour stimuli uninformative about reward association. **d,** Example animal performing the correlation discrimination task at different frequencies. Performance is calculated over a 100 trial sliding window. Insets: Trial maps before and after introduction of control valves (n = 12 trials pre-, n = 12 trials post-new valve introduction, new valve introduction indicated by black vertical dotted line, see also **Supp Fig 2.1**) **e,** Performance of 4 representative animals performing correlation discrimination where stimulus pulse frequency is randomised from trial to trial. Performance binned by frequency within approximately a half octave range across trials. **f,** Group performance in the random frequency experiment (black: standard training, orange: full scramble control). Performance binned by frequency within approximately a half octave range and across animals within each group (n = 33 training mice, n = 5 control mice).

Consistent with the fact that mice can detect temporally fluctuating odours, we find neurons that differentially respond to correlated and anti-correlated odours already in the output of the olfactory bulb using 2p imaging (**Fig 3a-d, Supp Fig 3.1**). Using linear classifier analysis (Cury and Uchida, 2010), we find that 10s of randomly chosen mitral/tufted cells are sufficient to guarantee the distinction of 20 Hz correlated and anti-correlated stimuli on a trial by trial basis (**Fig 3d,e**).

**Figure 3:**
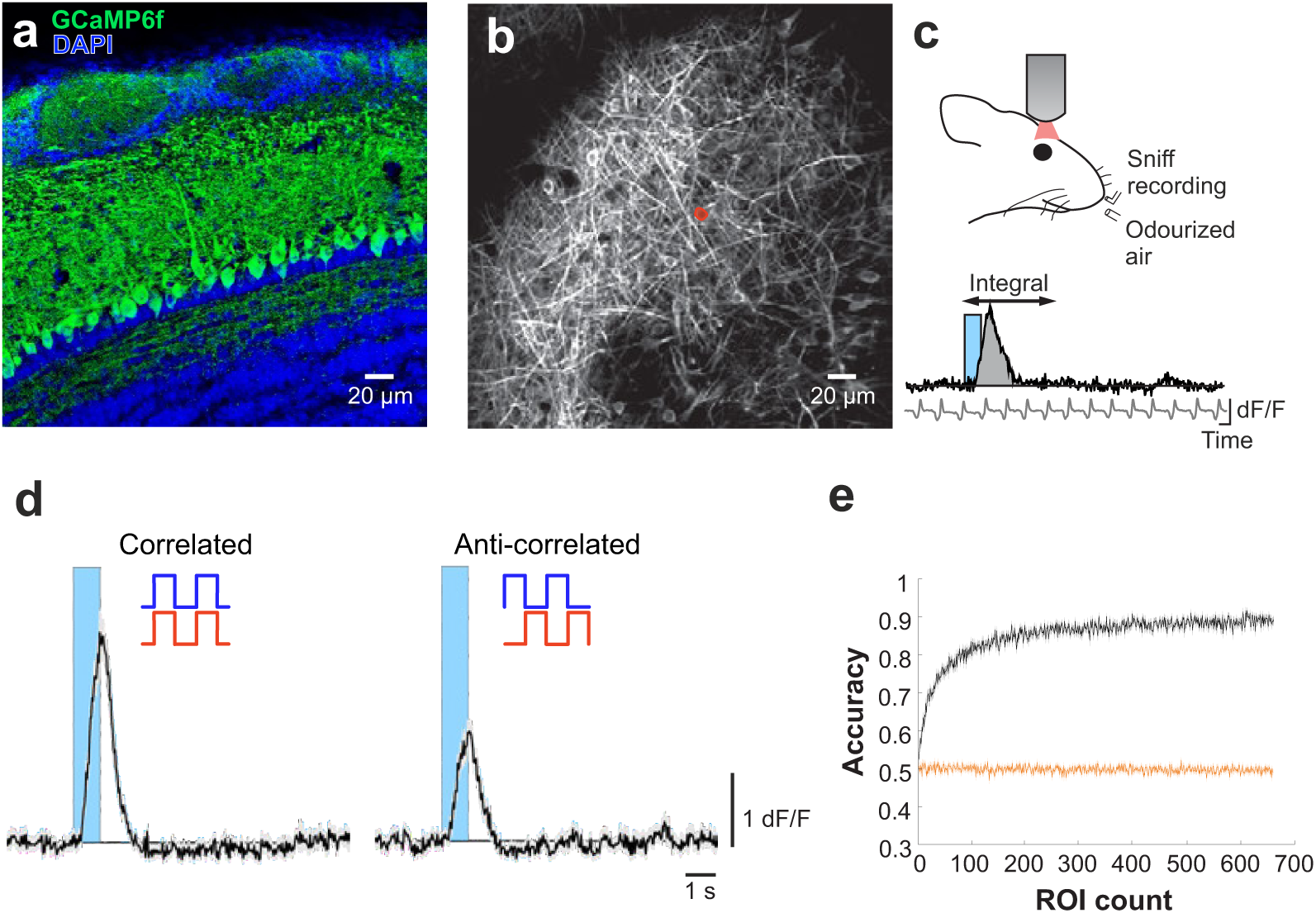
Correlation structure is encoded by olfactory bulb output neurons. **a,** Coronal section of the olfactory bulb showing GCaMP6f (green) expressed in projection neurons under the Tbet promoter. **b,** GCaMP6f fluorescence from mitral and tufted cells of the dorsal olfactory bulb imaged with a two-photon microscope through a cranial window (maximum projection of 8000 frames, red area marked is ROI that generated the response in **(d)**. Scale bars: 20 µm. **c,** Top: Schematic of the recording approach. Two-photon imaging of the dorsal olfactory bulb was performed while delivering correlated or anti-correlated odour stimuli. Respiration rate was simultaneously measured with a flow sensor placed close to the nostril contralateral to the bulb hemisphere being recorded. Bottom: integral of fluorescence measurements in a 5 s window after stimulus onset, including the 1 s stimulus, were used for further analysis. The respiration trace was used to align all responses to the first inhalation after stimulus onset. **d,** Example traces of the ROI marked in **(b)** in response to correlated (left) and anti-correlated (right) stimulation at 20 Hz (mean of 24 trials+/−SEM). **e,** Linear classifier accuracy over all ROIs when trained on all correlated vs. anti-correlated stimuli at 20 Hz (n = 661, mean +/− SEM; black: correlated vs. anti-correlated, orange: shuffle control).

Here, we have shown that mammals can detect temporal features of odour stimuli at frequencies of up to 40 Hz. This surprisingly high frequency is consistent with recent findings that the olfactory bulb circuitry not only enables highly precise odour responses (Cury and Uchida, 2010; Shusterman et al., 2011) but also enables detection of optogenetically evoked inputs with a precision of 10-30 ms (Rebello et al., 2014; Smear et al., 2011). While behavioural and physiological responses to precisely timed odour stimuli have been observed in insects (Brown et al., 2005; Geffen et al., 2009; Nagel et al., 2015; Riffell et al., 2014; Szyszka et al., 2014; Vickers et al., 2001), in mammals the complex structure of the nasal cavity was generally thought to “wash-out” any temporal structure of the incoming odour plume ((Kepecs et al., 2006) see however (Gupta et al., 2015)). Our results show that on the contrary mice can readily make use of information in odour stimuli fluctuating at frequencies of up to 40 Hz.

What could such high bandwidth be useful for? We have shown that odour sources even in close proximity differ in the temporal correlation structure. Thus, the behavioural ability to detect whether odorants are temporally correlated could allow mice to perform source separation segmentation, solving the “olfactory cocktail party problem” (Hopfield, 1991; Rokni et al., 2014) purely using temporal structure, without requiring the presence of unique chemicals in individual sources or prior knowledge of odours or components of odours (Grabska-Barwinska et al., 2017). Furthermore, extracting temporal features from odour fluctuations could allow for behavioural detection of distance or direction of an odour source (Celani et al., 2014; Justus et al., 2002; Mafraneto and Carde, 1994; Moore and Atema, 1988; Schmuker et al., 2016).

How could this temporal information be extracted? While insects are able to detect the simultaneity of onset of two odours (Baker et al., 1998; Stierle et al., 2013; Szyszka et al., 2012), we find that it is unlikely that this is the sole or even dominant means that mice use to detect correlation (**Supp Fig 2.4**). Similarly, mice do not show adjustment of sniff strategies in the behavioural experiment (**Supp Fig 2.5, Supp Video 2-3**). Already OSNs might be tuned to specific temporal features as shown in crustaceans (Park et al., 2014). While individual mammalian OSNs are thought to be quite slow and unreliable (Duchamp-Viret et al., 1999), the large convergence of OSN axons can provide a substrate to create the needed high temporal bandwidth (Abeles, 2004). Either intrinsic cellular biophysics (Hausser et al., 2000), local interneurons or long-range lateral inhibition (Fukunaga et al., 2014; Hopfield, 1991) might in turn permit the extraction of temporal correlation within the olfactory bulb circuitry.

The turbulence of odour plumes has often been viewed as a source of noise for mammals. Our finding that mice can extract temporal information from odour stimuli at a bandwidth of up to 40 Hz opens up a new perspective on how mice could make use of natural turbulence in order to obtain information about their spatial environment, providing new challenges to uncover what algorithms and computations the mammalian olfactory system might implement.

## Acknowledgements

We thank the animal facilities at National Institute for Medical Research and the Francis Crick Institute for animal care and technical assistance. We thank the mechanical and electronic workshops in Heidelberg (N. Neef, K. Schmidt, M. Lukat, R. Roedel, C. Kieser) and London (A. Ling, A. Hurst, M. Stopps) for excellent support during development and construction, the Aurora Scientific team for helpful suggestions for adapting the miniPID, T. Margrie for discussion and R. Jordan, C. Marin, J. Harris, F. Iacaruso, J. Kohl, and A. Fleischmann for comments on earlier versions of the manuscript. This work was supported by the Francis Crick Institute which receives its core funding from Cancer Research UK (FC001153), the UK Medical Research Council (FC001153), and the Wellcome Trust (FC001153); by the UK Medical Research Council (grant reference MC_UP_1202/5); a Wellcome Trust Investigator grant to AS (110174/Z/15/Z), and a DFG postdoctoral fellowship to TA.

## Methods

### Animals

All animal procedures performed in this study were approved by the UK government (Home Office) and by Institutional Animal Welfare Ethical Review Panel. All mice used for behavioural experiments were C57/Bl6 males.

### Reagents

All odours were obtained in their pure form from Sigma-Aldrich, St. Louis MO, USA. Unless otherwise specified, odours were diluted 1/5 with mineral oil in 15 ml glass vials (27160-U, Sigma-Aldrich, St. Louis MO, USA).

### High-speed odour delivery device

The odour delivery device was based on a modular design of four separate odour channels, and consisted of an odour manifold for odour storage, a valve manifold for control of odour release and hardware for controlling and directing air flow through the system. The odour manifold was a 12.2×3.2×1.5 cm stainless steel block with 4 milled circular indentations (1 cm radius). Within each of these indentations was a threaded through-hole for installation of an input flow controller (AS1211F-M5-04, SMC, Tokyo, Japan) and an output filter (INMX0350000A, The Lee Company, Westbrook CT, USA). For each inset, the cap of a 15 ml glass vial (27160-U, Sigma-Aldrich, St. Louis MO, USA) with the centre removed was pushed in and sealed with epoxy resin (Araldite Rapid, Huntsman Advanced Materials, Basel, Switzerland). This meant that glass vials could be screwed in and out of the insets.

Solenoid valves typically limit high-fidelity odour stimulation resulting in odour rise times of several 10s of milliseconds under optimal conditions (Raiser et al., 2017). We thus employed highspeed micro-dispense valves with custom electronics for pulse-width modulation to maximize bandwidth: 4 VHS valves (INKX0514750A, The Lee Company, Westbrook CT, USA) were installed in a 4-position manifold (INMA0601340B, The Lee Company, Westbrook CT, USA) with standard mounting ports (IKTX0322170A, The Lee Company, Westbrook CT, USA). Each valve was connected to a corresponding odour position in the odour manifold with 10 cm Teflon tubing (TUTC3216905L, The Lee Company, Westbrook CT, USA). Each valve was controlled by digital commands via a spike-and-hold driver. Each digital pulse delivered to the spike-and-hold driver delivered a 0.5 ms, 24 V pulse to the valve (to open it), followed by a 3.3 V holding pulse lasting the rest of the duration of the digital pulse. This spike-and-hold input allowed for fast cycling of the valve without switching between 0 and 24 V at high frequencies to prevent from overheating the valve. Each valve was controlled by an individual spike-and-hold driver. Up to 4 drivers could be controlled and powered with a custom-made PSU consisting of a 24 V power input and a linear regulator to split the voltages into a 24 V and 3.3 V line, as well as control inputs taking digital signal input and routing it to the appropriate valve. Pulse profiles for calibration and stimulus production were generated with custom Python software (PyPulse, PulseBoy; github.com/RoboDoig) allowing to define pulse parameters across multiple valves using a GUI.

To generate air flow through the olfactometer, a pressurised air source was connected to a filter (AME250C-F02, SMC, Tokyo, Japan) and demister (AMF250CF02, SMC, Tokyo, Japan) and then split into two separate lines, the input line and carrier line. Both lines were then connected to a pressure regulator (AR20-F01BG-8, SMC, Tokyo, Japan) and flow controller (FR2A13BVBN, Brooks Instrument, Hatfield PA, USA). The main line was then connected to the input of the valve manifold. The input line was split into 4 separate lines and connected to the input flow controllers (set to 0.25 L/min) on each odour position of the odour manifold. The output of the valve manifold was fitted with MINSTAC tubing (TUTC3216905L, The Lee Company, Westbrook CT, USA). Where the design was scaled up (e.g. to include 8 odour positions) the valve manifold outputs were connected and consolidated to a single output with 3-way connectors (QSMY-6-4, Festo, Esslingen am Neckar, Germany). Shape and reliability of odour pulses depended strongly on low volume headspace and low pressure levels (0.05 MPa).

#### Odour characterisation

Signal fidelities were calculated by first subtracting the average amplitude of troughs from the average amplitude of peaks during a pulse train and then subsequently dividing this peak-to-trough value by the difference of average peak amplitude subtracted by baseline amplitude. This results in a value between 0 and 1, with 1 being perfectly modulated odour pulses.

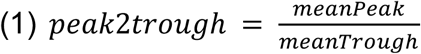

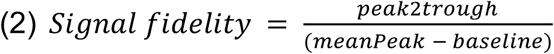

### Behaviour

#### Automatic operant conditioning of cohorts of mice (AutonoMouse)

In AutonoMouse, groups of mice (up to 25) implanted with an RFID chip are housed in a common home cage (Fig 2a, for detailed description see Erskine et al., 2019). Within the common home cage of AutonoMouse, mice have free access to food, social interaction and environmental enrichment. Water is not freely available in the system, but can be gained at any time by completion of an operant conditioning go/no-go task. To access these behavioural tasks, mice must leave the home cage and enter a behavioural area. This behavioural area contains the odour port and a lick port through which water rewards can be released. The lick port is also connected to a lick sensor which registers the animal’s response (its lick rate) in response to the task stimuli. As animals can only gain their daily water intake by completing behavioural tasks, mice are motivated to complete long sequences of trials without manual water restriction.

#### Training on temporally structured odours

We aimed to probe whether mice could perceive a particular temporal feature of naturally occurring odour signals: temporal correlations between odour signals. In particular we aimed to investigate this question with the simplest possible case: whether mice could discriminate perfectly correlated from perfectly anti-correlated odour stimuli.

All tasks followed a standard go/no-go training paradigm. Animals were presented with two odours presented in either a correlated pattern or an anti-correlated pattern (Fig 2b, Supp Fig 2.1a-c). For roughly half of all animals, the correlated pattern was S+ (rewarded) and the anti-correlated pattern was S-(unrewarded); in the other half of the group this reward valence was reversed. All stimuli were 2 s long. A water reward could be gained by licking such that licking was detected for at least 10 % of the stimulus time during an S+ presentation. Licking for the same amount of time during S-presentation resulted in a timeout interval of 7 s. In all other response cases the inter-trial interval was 3 s and no water reward was delivered.

#### Stimulus structure

All anti-correlated and correlated stimuli on each trial followed a common pattern in their construction. Generally, wherever an odour position is inactivated a blank position should be activated to compensate for flow change. There should also be no consistent differences in the amount of odour or flow released during the stimulus between correlated and anti-correlated stimuli. The detailed algorithm for stimulus generation is as follows:

1. Choose whether the stimulus will be correlated or anti-correlated.
2. A set of 1-2 positions each for odour 1 and odour 2 and 2-3 positions for blank are randomly chosen from a pre-defined subset of 6 of the 8 total positions. For example, a valid combination could be odour 1 at position 1, 2; odour 2 at position 5; and blank at position 3 and 7. (see Fig 2c, Supp Fig 2.1b)
3. Create a guide pulse at the desired frequency (e.g. 2 Hz pulse with 50% duty) for all positions that follows the chosen stimulus structure.
4. The relative contributions of each position to the total stimulus are randomly generated. At each time point in the stimulus, only two position types should be active (e.g. odour 1 and blank for an anti-correlated stimulus) so the maximum contribution for any position type is 50% of the total release amount. Where two positions have been chosen for a position type, their relative contributions should add to 50%.
5. The guide pulses are pulse-width modulated according to the relative contributions of each position. PWM is at 500 Hz with some added jitter in the duty to avoid strong tone generation.

#### Task structure

Task frequency was randomised from trial to trial in a range between 2-81 Hz. The choice of frequency was with weighted probability divided into 3 frequency bands. E.g. this task could be arranged such that 2-20 Hz would be chosen with P = 0.6, 21-40 Hz with P = 0.3 and 41-81 Hz with P = 0.1. Within each of these frequency bands, the choice of individual task frequency was based on a uniform distribution.

#### Onset detection

For the onset detection experiments (Supp Fig 2.4) animals were trained to discriminate perfectly correlated (e.g. S+) from perfectly anti-correlated stimuli (e.g. S-) and probed with partially altered stimuli where the onset (first cycle) of the probe S+ stimuli was anti-correlated and probe S-stimuli where the onset (first cycle) was correlated. Performance during these probe trials is then compared to the average performance during training (*perf*_*train*_).

We calculated the expected average animal performance on the probe trials based on two models: Model 1 assumed the animals were taking any part of the stimulus into account equally when making a decision. Model 2 assumed that only the onset of the stimulus would contribute to discrimination. For Model 1 thus, a stimulus of frequency *f* (e.g. 10 Hz) that was sampled for *t*_*sample*_consisted of a “shifted” onset component of one cycle for S+ (1/*f*) and half a cycle for S-(0.5/*f*) corresponding to a fraction of *frac*_*onset*_= 1/*f*/*t*_*sample*_of the entire stimulus and a “normal” residual (*frac*_*res*_= 1-*frac*_*onset*_). Thus the predicted probe trial performance would be:

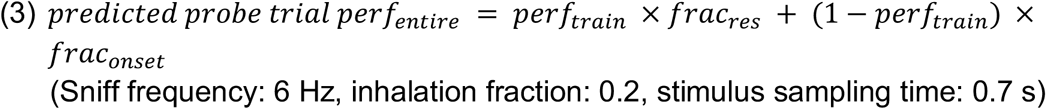

For Model 2, ignoring inhalation timing, the prediction would be that preference would be reversed (as onset correlations during probe trials are reversed). However, this ignores the fact that odour stimuli during the exhalation period might not be detected. Thus, to more accurately predict animals’ performance for Model 2, we assume that the part of the stimulus that is detected as the “onset” is the first odour pulse during an inhalation phase. During the probe trial, this will be the “inverted” first cycle if the stimulus begins either during the inhalation phase or at most 1/*f* before the inhalation (then inhalation would start during the inverted first cycle of the probe trial). The probability of this occurring is *perf*_*onset*_= (*dur*_*inh*_+1/*f*) / *dur*_*sinff*_with *dur*_*inh*_and *dur*_*sinff*_being inhalation and sniff duration respectively (provided *dur*_*inh*_+1/*f* < *dur*_*sinff*_). Predicted probe trial performance for an “onset only” model would thus be:

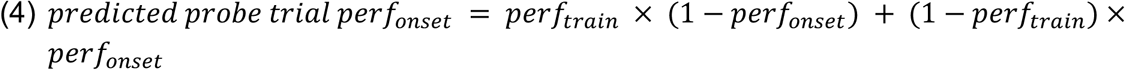

#### Controls

Control valves could be automatically added to the random frequency task. These tasks produced their stimuli based on a subset of 6 valves, control valves could be added automatically after a set period of trials to force the algorithm to produce stimuli from all 8 valves (see Fig 2c, switch control).

In switch controls, the experiment was halted and the positions of all valves and odour positions were shuffled in the olfactometer (see Fig 2c, switch control). The software definitions for valve positions were remapped to account for the change and the experiment was restarted.

A subgroup of animals was created in which the valve map was scrambled, as an ongoing control against animals learning extraneous variables in the task (see Fig 2c, scramble control). The valve map was scrambled in the following way: One blank to odour 1, one odour 2 to blank, one odour 1 to odour 2 and one odour 1 to blank.

#### Cohorts

The correlation discrimination experiment was performed in 2 separate experimental cohorts (group 1, n = 14; group 2, n = 25). Each cohort was organised into several subgroups which performed slight variations of the behavioural tasks in terms of reward valence and valves utilised, but with the same underlying task aim. Half of the animals in each subgroup were trained on correlated stimuli as the S+ rewarded condition, with the other half trained on anti-correlated as rewarded. Animals were further subdivided into groups which were trained on different subsets of valves as standard in the 8-channel olfactometer. For each cohort, mice were once assigned to each of these subgroups based on performance in a simple pure odour discrimination at the beginning of the experiment - group membership was randomised until no significant (ANOVA, Tukey-Kramer) differences in performance could be extracted between these subgroups on this task.

#### Data Analysis

AutonoMouse behavioural data was converted to MATLAB data format using the Conversion module of the Python autonomouse-control package (github.com/RoboDoig). All subsequent analysis was performed with custom-written MATLAB scripts unless otherwise specified.

All behavioural performance within a specified trial bin was calculated as a weighted average of S+ vs. S-performance:

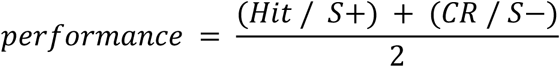

Where S+ is the total number of rewarded trials, S-is the total number of unrewarded trials, Hit is the total number of rewarded trials in which a lick response was detected, CR (correct rejection) is the total number of unrewarded trials in which no lick response was detected.

For random stimulus pulse frequency experiments (e.g. Fig. 2e,f) trials were binned approximately by half-octave for performance analysis. The exact intervals were Hz = [2, 3, 4, 5, 6:7, 8:10, 11:13, 14:17, 18:22, 23:29, 30:37, 38:48, 49:62, 63:81]. Reaction time (Supp Fig. 2.2) was calculated from S+ trials for each animal as the time to the first lick after stimulus onset.

Motion magnification of the respiration camera video recordings was performed with phase-based video motion processing with correction for large body movements based on MATLAB scripts by Wadhwa et al., 2013 (phaseAmplifyLargeMotions). Parameters for phase amplification were: blurring σ = 1, magnification α = 50, amplification in frequency band between 2-13 Hz. Following magnification, static ROIs for each video were selected in Bonsai (http://www.kampff-lab.org/bonsai/, Lopes et al., 2015) over the animal flank. An adaptive binary threshold was applied to the ROI to segment the animal body from the video background. Respiration rate was extracted as the total size of the ROI occupied by the body over time.

### In vivo two-photon imaging

#### Surgical and experimental procedures

Prior to surgery all utilised surfaces and apparatus were sterilised with 1% trigene. 12-20 week old Tbet-cre mice (Haddad et al., 2013) crossed with a GCaMP6f reporter line (Otazu et al., 2015) were anaesthetised using a mixture of fentanyl/midazolam/medetomidine (0.05 mg/kg, mg/kg, 0.5 mg/kg respectively). Depth of anaesthesia was monitored throughout the procedure by testing the toe-pinch reflex. The fur over the skull and at the base of the neck was shaved away and the skin cleaned with 1 % chlorhexidine scrub. Mice were then placed on a thermoregulator (DC Temperature Controller, FHC, ME USA) heat pad controlled by a temperature probe inserted rectally. While on the heat pad, the head of the animal was held in place with a set of ear bars. The scalp was incised and pulled away from the skull with four arterial clamps at each corner of the incision. A custom head-fixation implant was attached to the base of the skull with medical super glue (Vetbond, 3M, Maplewood MN, USA) such that its most anterior point rested approximately 0.5 mm posterior to the bregma line. Dental cement (Paladur, Heraeus Kulzer GmbH, Hanau, Germany; Simplex Rapid Liquid, Associated Dental Products Ltd., Swindon, UK) was then applied around the edges of the implant to ensure firm adhesion to the skull. A craniotomy over the left olfactory bulb (approximately 2 x 2 mm) was made with a dental drill (Success 40, Osada, Tokyo, Japan) and then immersed in ACSF (NaCl (125 mM), KCl (5 mM), HEPES (10 mM), pH adjusted to 7.4 with NaOH, MgSO4.7H2O (2 mM), CaCl2.2H2O (2 mM), glucose (10 mM)) before removing the skull with forceps. The dura was then peeled back using fine forceps. A layer of 2 % low-melt agarose diluted in ACSF was applied over the exposed brain surface before placing a glass window cut from a cover slip (borosilicate glass 1.0 thickness) using a diamond knife (Sigma-Aldrich) over the craniotomy. The edges of the window were then glued with medical super glue (Vetbond, 3M, Maplewood MN, USA) to the skull.

Following surgery, mice were placed in a custom head-fixation apparatus and transferred to a two-photon microscope rig along with the heat pad. The microscope (Scientifica Multiphoton VivoScope) was coupled with a MaiTai DeepSee laser (Spectra Physics, Santa Clara, CA) tuned to 940 nm (<50 mW average power on the sample) for imaging. Images (512 x 512 pixels) were acquired with a resonant scanner at a frame rate of 30 Hz using a 16x 0.8 NA water-immersion objective (Nikon). The output of a 4-channel version of the temporal olfactometer described above was adjusted to approximately 1 cm away from the ipsilateral nostril to the imaging window, and a flow sensor was placed to the contralateral nostril for continuous respiration recording.

#### Awake recordings

For implantation of the head-plate, mice were anaesthetized with isoflurane in 95% oxygen (5 % for induction, 1.5 % - 3 % for maintenance). Local (mepivacaine, 0.5 % s.c.) and general analgesics (carprofen 5 mg/kg s.c.) were applied immediately at the onset of surgery. After surgery, animals were allowed to recover for 7 days with access to wet diet and, after recovery, habituated to the head-fixed situation for at least 15 min on three consecutive days preceding the imaging experiment (**Supp Fig 3.1**).

#### Odour stimulation

Stimuli were generated from mixtures of physically mixed monomolecular odorants in order to ensure high probability of finding odour responsive cells in the dorsal olfactory bulb using custom Python Software (PulseBoy). Binary mixtures were diluted in mineral oil at the ratio of 1:5 and installed into the 4-channel olfactometer (15 ml per vial) along with two blank positions (15 ml mineral oil). Mix A: ethyl butyrate + 2-hexanone, mix B: isopentyl acetate + cineole. For all stimuli, odour valve offsets were compensated by opening a corresponding blank position valve to ensure no global flow changes occurred over the course of the stimulus. All stimuli were repeated 24 times with a 20 s inter-stimulus interval.

#### Analysis

Initial analysis was performed with custom scripts in Fiji. From an average of ∼8000 frames, ROIs around cell somata were manually selected and calcium transients were extracted and exported for further analysis in MATLAB. All traces were aligned to the first inhalation after odour onset. Calcium response integrals were calculated in a 5 s window from odour onset. To analyse how well odour responses predicted stimulus correlation on a trial-to-trial basis, we generated a linear discriminant classifier from the data set and analysed prediction accuracy. For the classifier, we performed 50% holdout validation, splitting the data randomly into a training set and test set with equal numbers of samples. We then performed linear discriminant analysis on the training data set to determine the best linear boundary between correlated vs. anti-correlated data. Classifier performance was then validated on the test data set. To determine the effect of number of ROIs used on classifier performance, we iteratively trained multiple classifiers on random subsets of ROIs with increasing numbers of ROIs within each set. For each ROI subset size, 100 classifiers were trained and the mean +/−SEM of their performance accuracy was calculated.

### Dual-energy fast photoionisation detection (defPID)

Two photoionisation detectors (200B miniPID, Aurora Scientific, Aurora ON, Canada) fitted with UV lamps of emission energy 10.6 eV (PID high) and 8.4 eV (PID low) were used to discriminate ethyl butyrate (EB, ionisation energy = 9.9 eV) from α-Terpinene (AT, ionisation energy = 7.6 eV) or ethyl valerate (EV, ionisation energy = 10.0 eV) from tripropyl amine (TA, ionisation energy = 7.2 eV). The PID inlets were connected with a 3-way connector to detect incoming odours by both PIDs simultaneously from a common point. PID heads were held on lab stands with the PID inlet at approximately 4 cm above ground level.

#### Odour delivery

Odours were held in ceramic crucibles (5 cm diameter, 6 ml volume) covered in an air-tight fashion using glass lids. Odours were released for 5 s with an inter-trial interval of 15 s by arduino-based robots programmed to lift the lids from the crucibles using a servo motor (TowerPro SG-5010, Adafruit, UK). PID recordings and robot movements were remotely controlled and synchronized from a computer. Experiments were carried out in a large open space, both indoors and outdoors.

Outdoors setup: PIDs and odour delivery system as described above were used to record for multiple trials in different conditions on a day with low wind (∼8-12 mph, based on BBC weather).

Indoors setup: A digitally controlled fan (2214F/2TDH0, ebm-papst, Chelmsford, UK) was placed at a distance of 440 cm facing the PID inlet. An exhaust line was situated behind the PID inlet to ensure the direction of air from the fan towards the PID inlet. During a recording, the fan was set to maximum speed such that it pushed approximately 552 cf/min (cubic feet per minute) of air towards the PID inlet. A 25×25×25 cm Thermocool box was placed 200 cm downwind of the fan acting as an obstacle to air movement, breaking up any laminar air flow and ensuring turbulent air movement at the PID inlet.

#### Recording conditions

6 ml of the desired odour(s) were filled in two crucibles and placed in different locations based on the experimental conditions as described below:

1. Low energy only: The low-energy odour (AT or TA) was placed 40 cm (radial distance) away from the PID inlet, and displaced either 25 cm left or 25 cm right of the midline (the midline in this context is the line between the PID inlet and the centre of the fan). The odour source was alternated between left and right positioning relative to the midline an equal number of times to remove any possible bias from positioning in the air stream. The purpose of this recording condition was to generate data to calculate the linear transformation from the low energy signal to the high energy signal (Supp Fig1.1c,d).
2. Mix: 3 ml EB + 3 ml AT (or 3 ml EV + 3 ml TA) was pipetted in one crucible and placed either 25 cm left or 25 cm right of the midline at radial distances of 20 cm, 40 cm and 60 cm. The purpose of this recording condition was to determine how the temporal structure of individual odours in a plume behaved when the odours emitted from the same source.
3. Separate: 3 ml EB and 3 ml of AT (or 3 ml EV + 3 ml TA) were individually pipetted in two different crucibles and placed at a radial distance of 40 cm from the PID inlet. For the 50 cm apart condition, one odour source was placed 25 cm left of the midline while the other 25 cm on the right of the midline and vice-versa (equal number of trials for both cases) separating the odour sources by 50 cm. This procedure was repeated for lateral distances of 30 cm and 10 cm. The ‘50 cm apart’ case was repeated for radial distances of 20 cm and 60 cm. The purpose of this recording condition was to determine how the temporal structure of individual odours in a plume behaved when the odours emitted from separated sources but were still free to mix in air.

#### Analysis

Decomposition procedure: The low energy odour (AT) was recorded using both PIDs. Assuming a linear relation between the recorded signals from the 2 PIDs, we plotted the recorded events with a linear regression fit (Supp Fig 1.1c) and calculated slope and R^2^ value of the fit. The scaling factor (6.82 +/−SD 0.356) was calculated as the average slope of all linear fits for R^2^ ≥ 0.9.

The PID low traces were multiplied by this scaling factor which was termed as the estimated low energy odour (Supp Fig 1.1e). The estimated high energy odour was calculated by subtracting the estimated low energy odour from the PID high traces.

Correlation calculation: Custom written scripts in MATLAB (Mathworks, USA) were used to calculate the correlation coefficient between the estimated low energy odour and the estimated high energy odour for all conditions. Box plots were obtained from these values using Igor Pro 6 (WaveMetrics, USA).

---

**Supplementary Figure 1.1:**
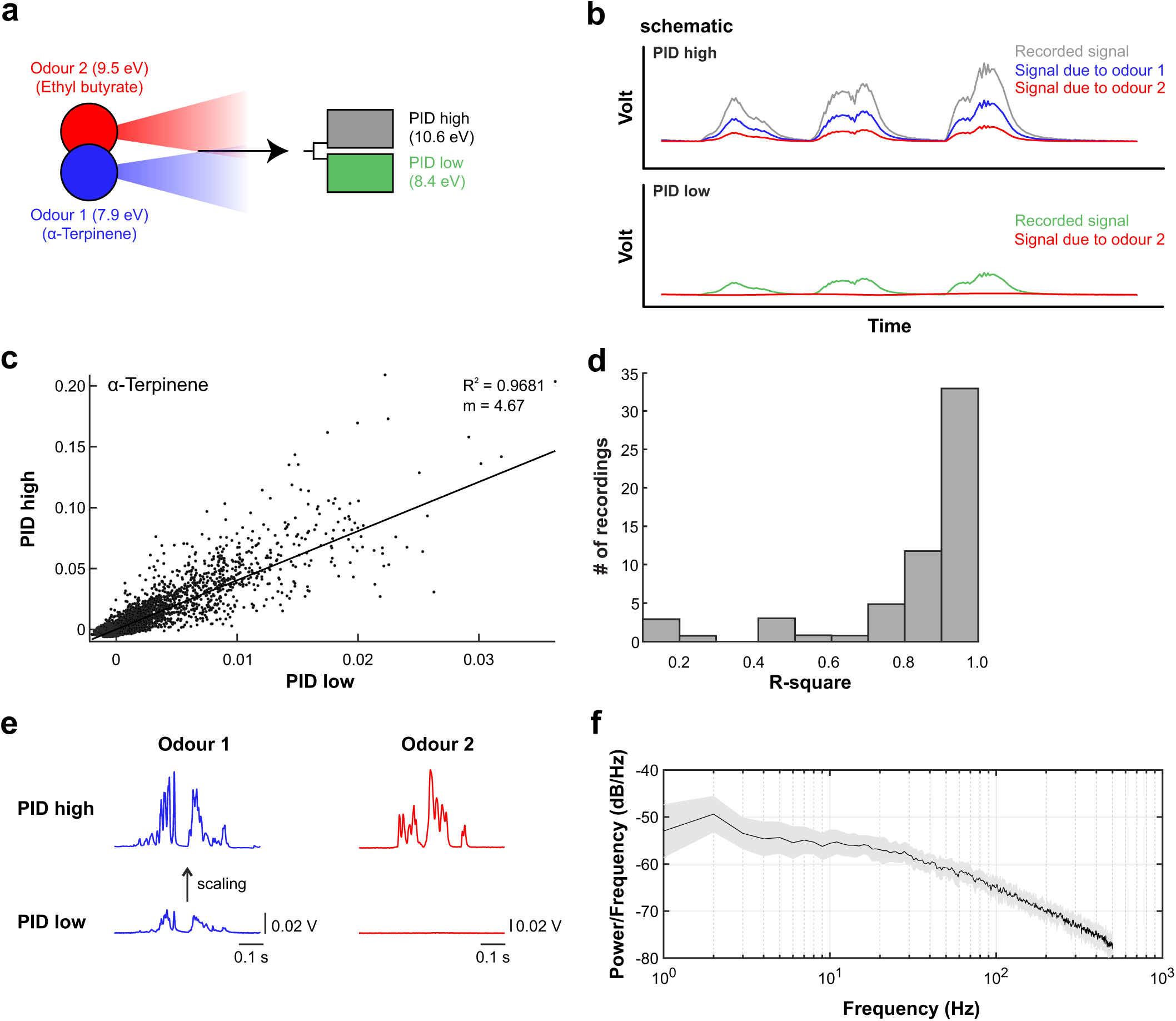
Dual-energy fast photoionisation detection (defPID). **a,** Schematic of the dual-energy fast photoionisation detection method. Two odours are recorded simultaneously by two PIDs with different ionising energies (different wavelength UV light sources). The odours are chosen such that one odour (odour 2 (Ethyl butyrate), 9.5 eV) has an ionisation energy greater than the low energy PID bulb, but less than the high energy PID bulb, thus only being detectable by the high energy PID. The other odour (odour 1 (α-Terpinene), 7.9 eV) is chosen such that its ionisation energy is lower than both PID bulbs (detectable by both PIDs, see also **(e). b,** Method of decomposing odour signals. Top panel: high energy PID signal (grey: recorded signal, blue: calculated signal due to odour 1, red: calculated signal due to odour 2). Bottom panel: low energy PID signal (green: recorded signal, red: calculated signal due to odour 2). **c,** Single data points of the PID signal evoked by α-Terpinene in the two PIDs. The slope of the linear fit serves as a scaling factor to map the low energy PID to the high energy PID signal. **d,** Histogram of R-square values of all dual-PID α-Terpinene recordings to define the scaling factor (n = 59). **e,** Summary of signal combinations for defPID recordings. **f,** Averaged power spectrum of odour plumes recorded under four different experimental conditions (α-Terpinene mix, Ethyl butyrate mix, α-Terpinene and Ethyl butyrate, radially separated by 40 cm).

**Supplementary Figure 1.2:**
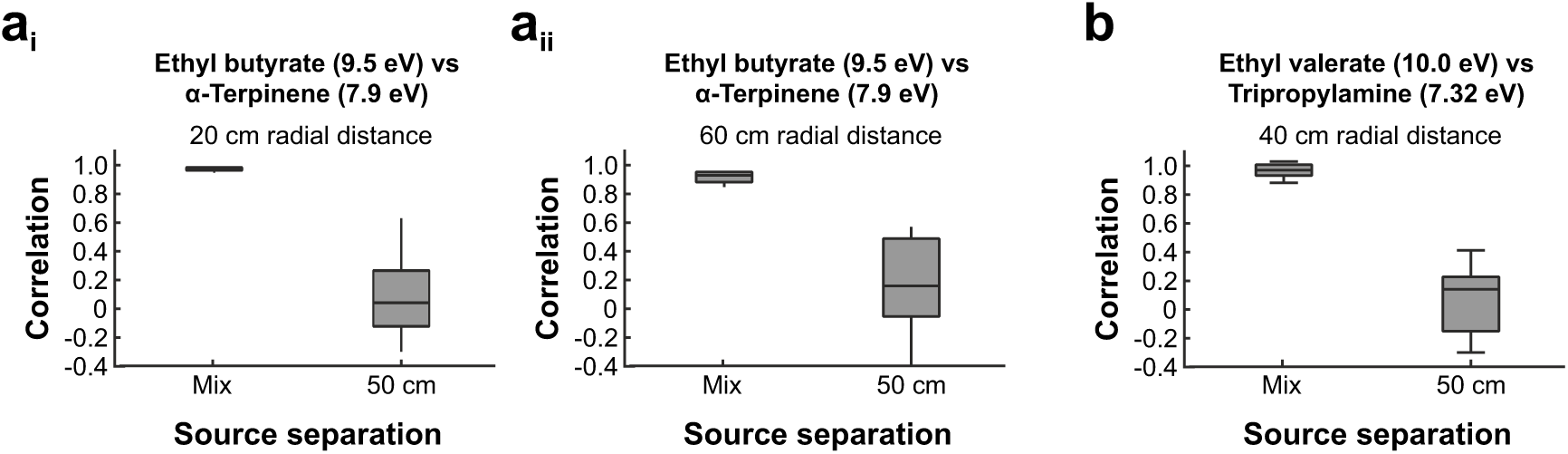
defPID measurements with different radial distances and with a different odour pair. **a,** Average odour signal correlation over all recordings (mean +/−SD) of α-Terpinene and Ethyl butyrate at 20 cm (n = 70) **(ai)** and 60 cm (n = 57-83) **(aii)** radial distance. **b,** Average correlation over all recordings (mean +/−SD) of Ethyl valerate and Tripropylamine at 40 cm radial distance (n = 25-27).

**Supplementary Figure 1.3:**
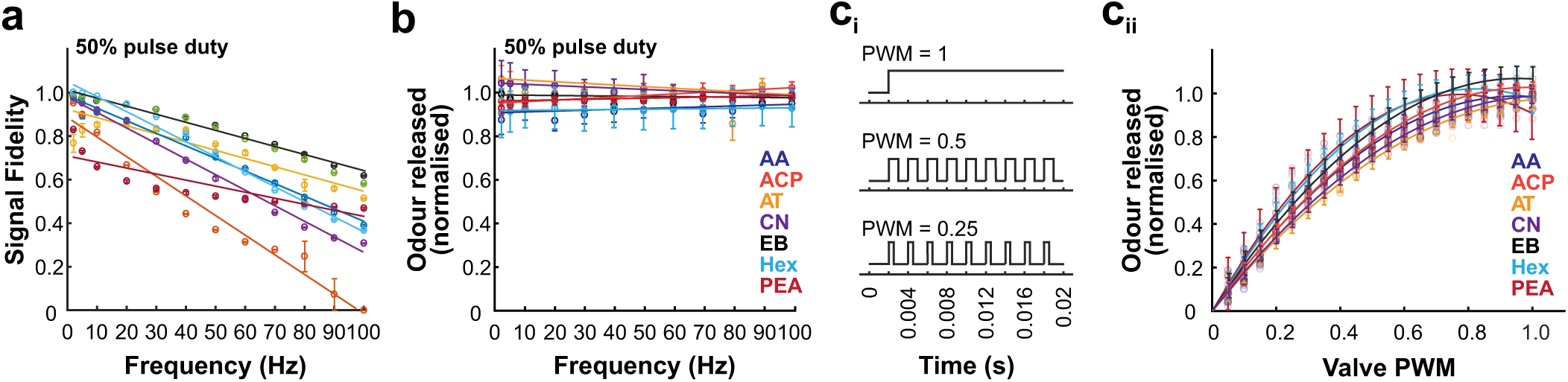
Characterization of odourants presented with high-speed odour delivery device. **a,** Calculated signal fidelities and **b,** amount of released odour for seven different odours pulsed for 2 s over a frequency range of 2 to 100 Hz at 50% pulse duty (n = 5 repeats for each condition, mean +/−SEM). **c,** Pulse width modulation control. Schematic of the pulse-width modulation (PWM) method **(ci)**. For any period of odour release, the maximum amount of odour release is achieved by holding the valve consistently open (top). The amount of odour released can be reduced by cycling the valve at a high frequency (here 500 Hz) with a different level of pulse-width modulation (middle and bottom panel). Odours were released over a 2 s period with different pulse width modulation duties at 500 Hz (**cii**, n = 5 repeats for each condition). The resulting released odour amount is normalised to the maximum release (PWM = 1). Odours are: AA (Isoamyl acetate), ACP (Acetophenone), AT (α-Terpinene), CN (Cineol), EB (Ethyl butyrate), Hex (2-hexanone), PEA(Phenylethyl alcohol).

**Supplementary Figure 1.4:**
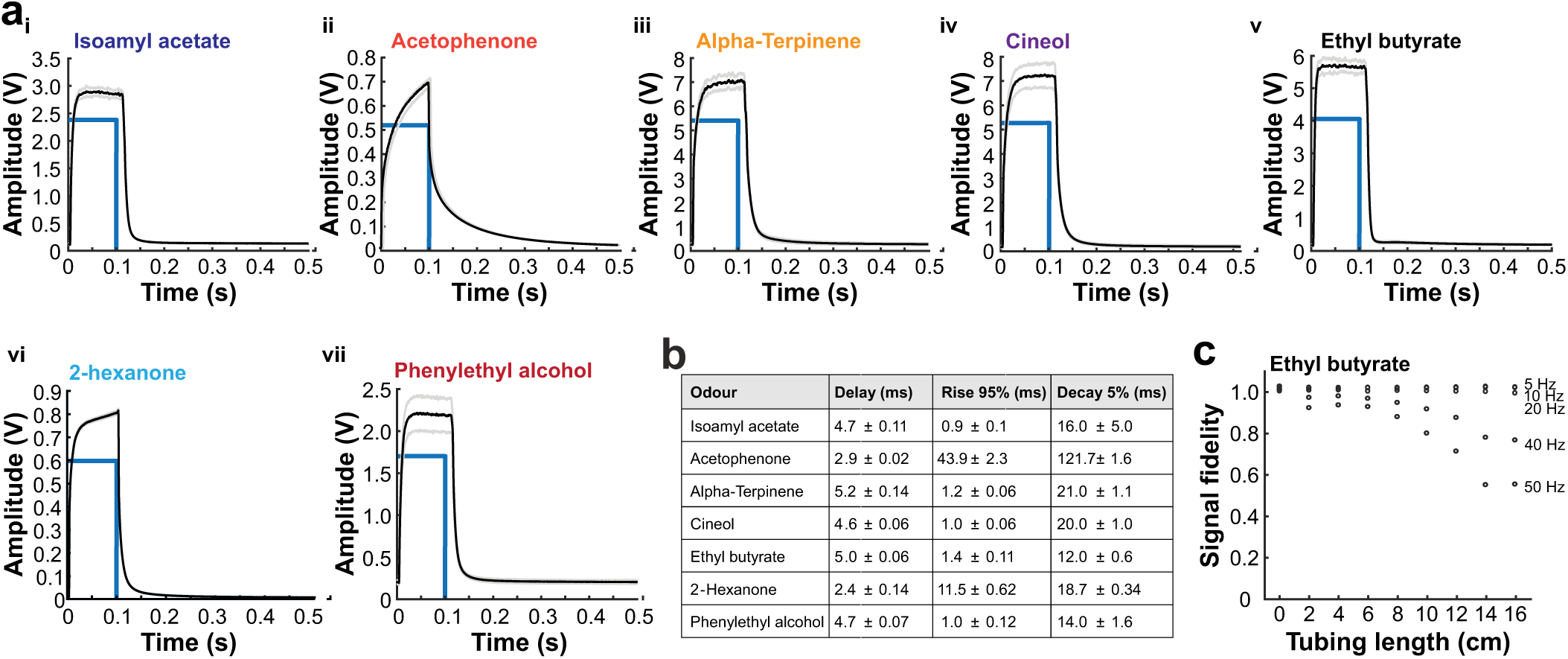
Characterisation of intrinsic odour properties. **a,** Average PID signal of single 100 ms pulses (pulse indicated in blue) for seven different odours (n = 60 pulses for each odour, mean +/−SEM, odours as in Supp Fig 1.3). **b,** Summary table: delay (time from start of the odour pulse to 5% of maximum signal amplitude), rise (time from 5% to 95% of maximum signal amplitude), decay (time for the signal to decay back to 5% of maximum amplitude after odour pulse). **c,** Effect of the length of tubing attached to the valve manifold on signal fidelity at different pulse frequencies for Ethyl butyrate pulsed for 2 s at 50% pulse duty.

**Supplementary Figure 2.1:**
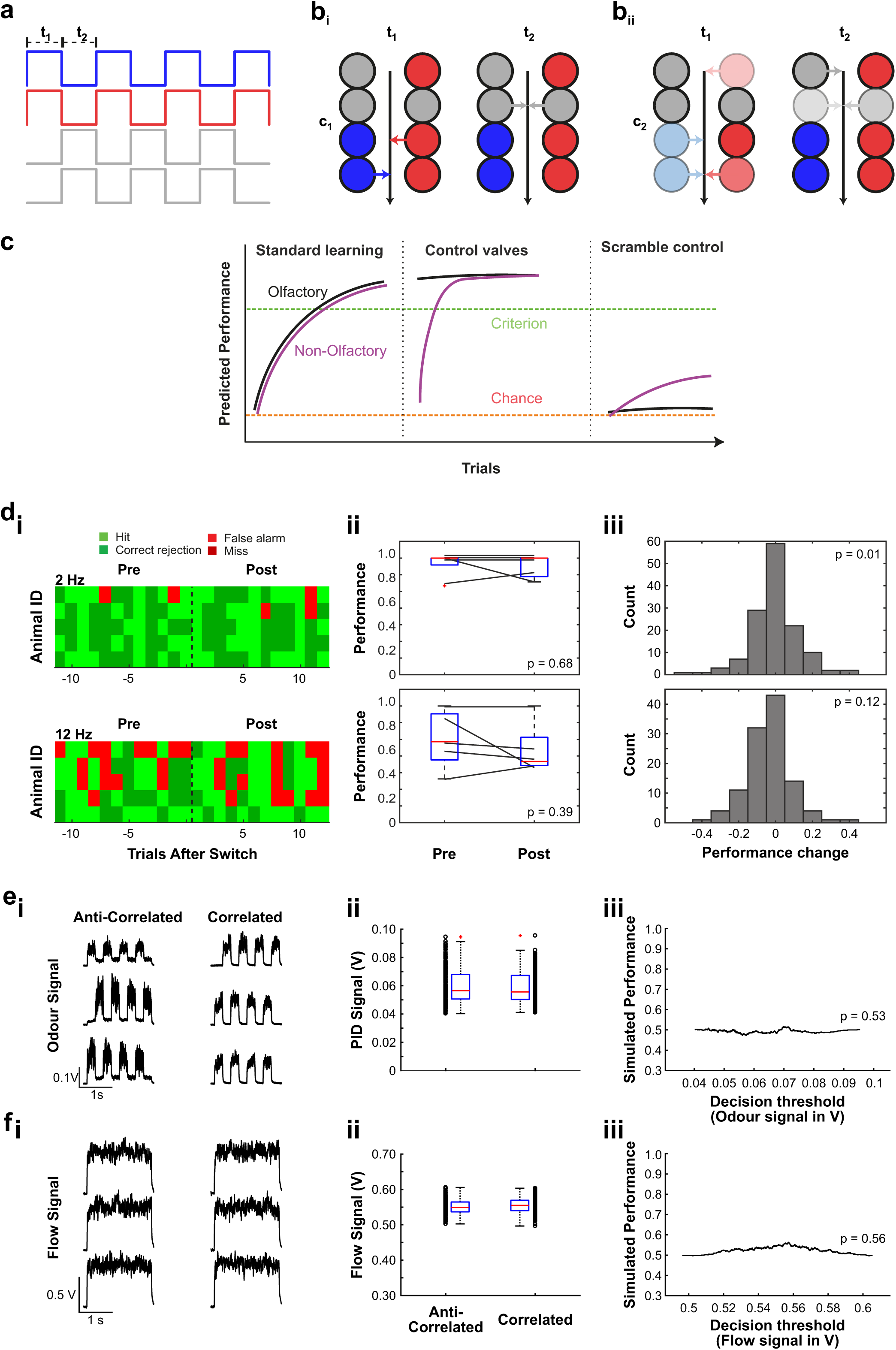
Stimulus and experimental design. **a,** Detailed schematic of stimulus production; odour presentation (blue = odour 1, red = odour 2) is always offset by clean air (grey) valves at the same flow levels, to ensure that total flow during the stimulus is constant. **b,** Schematic of the use of valve subsets to produce the desired stimulus. t1 and t2 represent valve openings at the corresponding time points shown in **(a)**. c1 **(bi)** and c2 **(bii)** represent two possible configurations that could be used to produce the same resulting stimulus at the two time points. Opacity in the colours represents total concentration contribution to the resulting stimulus at the time point. For example, to produce the dual odour pulse at t1, configuration c1 can be used in which odour 1 (blue) is generated by 50% opening of two valves, with odour 2 (red) produced by 70% / 30% opening of two valves respectively. **c,** Predicted performance for animals in the case that they use solely olfactory temporal correlations (black) and in the case that they use extraneous non-olfactory cues or non-intended olfactory cues (e.g. contaminations) (violet). Note that when switching stimulus preparations to a new set of valves, such non-intended cues would not provide any information about stimulus-reward association, thus animals performance would transiently drop back to chance **d,** Trial map of 5 representative animals during 2 Hz and 12 Hz correlation discrimination tasks before and after introduction of control valves (n = 12 trials pre-, n = 12 trials post-new valve introduction, new valve introduction indicated by black vertical dotted line. Each row corresponds to an animal, each column within the row represents a trial. Light green: hit, dark green: correct rejection, light red: false alarm, dark red: miss **(di)**. Boxplots of mean performance for selected animals pre-and post-control **(dii)**. Summary histograms of performance change for all animals during switch control tests **(diii)**. In no case was post-control performance significantly changed from pre-control performance suggesting that all discrimination performance was based on intended olfactory cues only. **e,** Example traces of odour signal (ethyl butyrate, amyl acetate, PID recorded) during anti-correlated trials (left) and correlated trials (right) **(ei)**. Mean odour signal for all recorded anti-correlated and correlated stimuli **(eii)** (total anti-correlated: n = 300, total correlated: n = 300. Within each subgroup: 30 trials each at 2 Hz, 5 Hz, 10 Hz, 20 Hz, 50 Hz). Box plots show median (red line), 25th to 75th percentile (blue box), most extreme points not considered outliers (black dashed line) and outliers (red +). Simulated maximum performance based on differences in mean odour signal **(eiii)**. Simulated performance was calculated as the fraction of trials correctly identified as correlated / anti-correlated based on a decision threshold set at some level between the minimum and maximum mean signal. Simulated performance was calculated for multiple decision thresholds, increasing the decision threshold from minimum odour signal to maximum odour signal in steps of 1/5000th of the range between minimum and maximum. **f,** Example traces of air flow recordings for the same trials described in **(e)**.

**Supplementary Figure 2.2:**
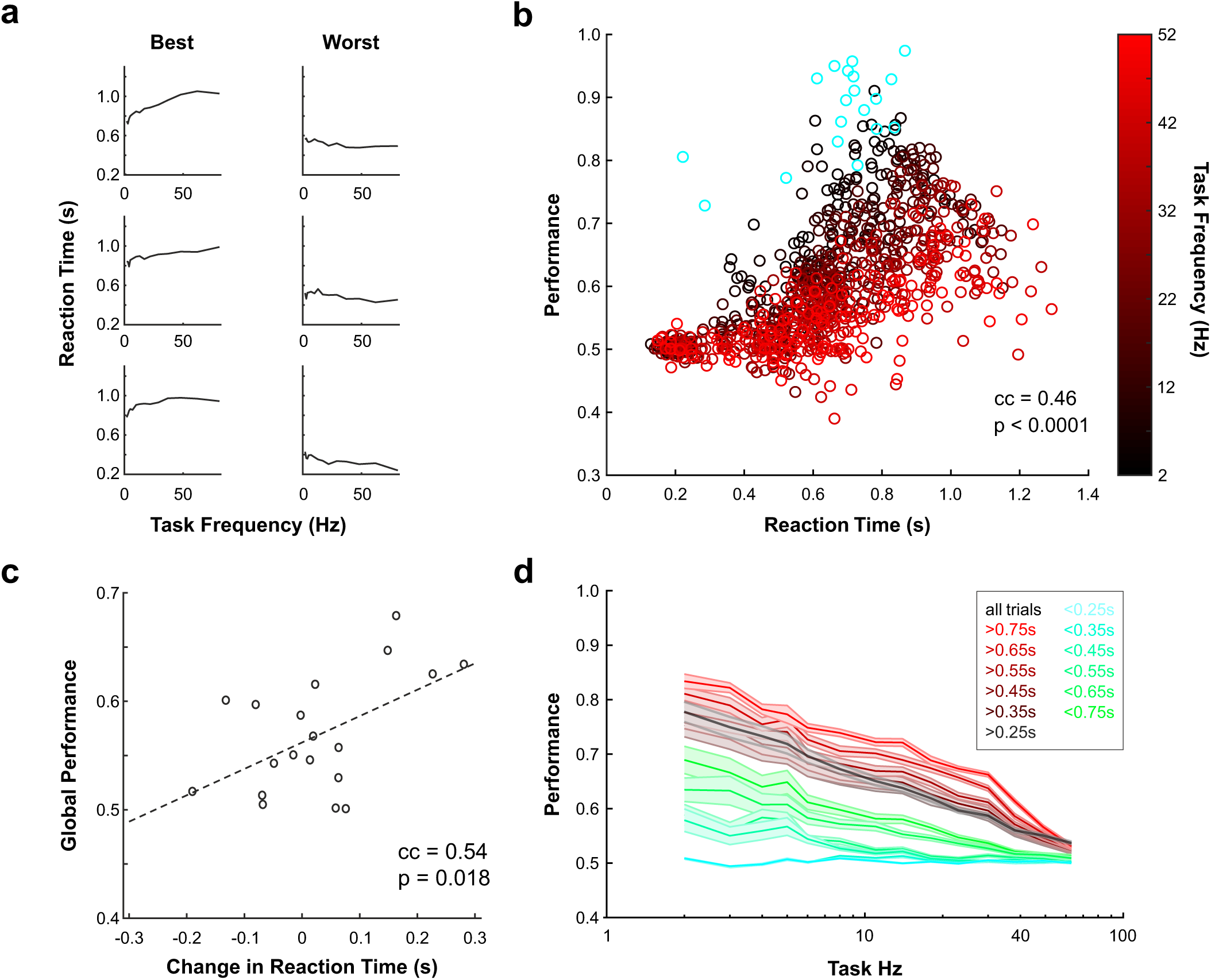
Reaction times. **a,** Mean reaction time (time from stimulus onset to first lick in S+ trials) are plotted as a function of stimulus pulse frequency (from random frequency experiment in **Fig. 2e,f**) for the three animals with the best global performance (mean performance across all trials) and the worst global performance. Better performing animals tend to increase their reaction time as stimulus pulse frequency increases. **b,** Scatter plot of mean performance vs. mean reaction time for each animal and stimulus pulse frequency condition. Points are colour coded according to stimulus pulse frequency. Blue points are for animals performing a simple discrimination between two pure monomolecular odorants. Performance was significantly positively correlated to reaction time, suggesting that mice that sampled a greater portion of the stimulus made more accurate decisions about its correlation structure. **c,** For each animal, the change in reaction time (defined as mean reaction time at the lowest stimulus pulse frequency subtracted from the mean reaction time at the highest stimulus pulse frequency) is plotted against the global performance as defined in **(a)**. A positive change in reaction time indicates that an animal increased its stimulus sampling time for higher stimulus pulse frequencies. There was a positive and significant correlation between change in reaction time and global performance, suggesting that animals that sample the stimulus for longer at higher stimulus pulse frequencies were better able to discriminate odour temporal correlation structure. **d,** Performance is plotted as in **(Fig. 2f)** on the random frequency experiment, but only animals with reaction times above or below a certain threshold (colour code) are included in the analysis. Where only longer reaction times are considered, global performance is higher than the case where only shorter reaction times are included, suggesting that longer stimulus sampling improves discrimination of odour correlation structure across all stimulus pulse frequencies.

**Supplementary Figure 2.3:**
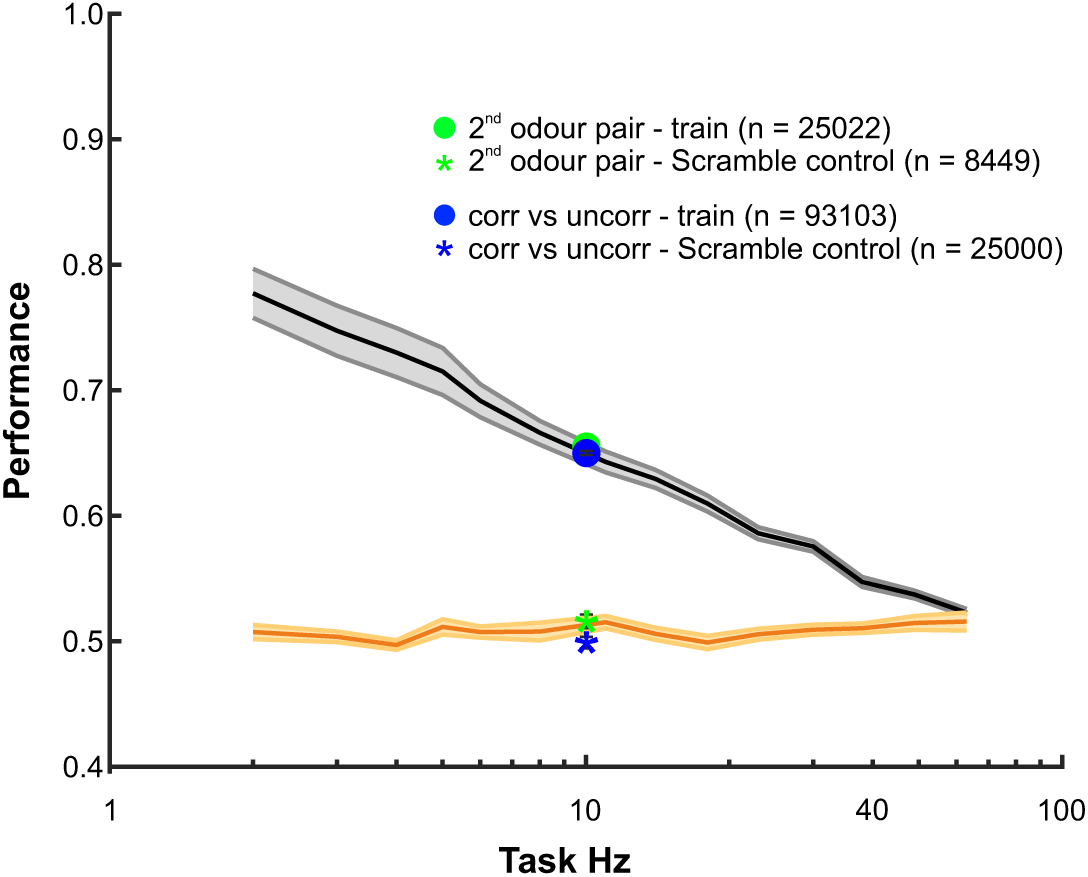
Discrimination performance for second odour pair and correlated versus uncorrelated stimuli. Animals show similar average performance when asked to discriminate the correlation structure of a different odour pair (Acetophenone vs. Cineol) at 10 Hz (green dot, n = 19, mean performance = 0.6558 +/−0.0026; mean performance = 0.5165 +/−0.0048 when stimulus identity was scrambled). In addition to testing the discrimination ability of correlated and anti-correlated odour pulses, animals were probed to discriminate correlated from uncorrelated odour pulses at 10 Hz. Average performance is still significantly above chance when pulse timing of one odour is randomised until correlation of both pulses is 0 (blue dot, n = 19, mean performance = 0.6506 +/−0.0016; mean performance = 0.4997 +/−0.0032 when stimulus identity was scrambled). Plots from **(Fig 2f)** for test animals (black) and scramble control animals (orange) shown for comparison.

**Supplementary Figure 2.4:**
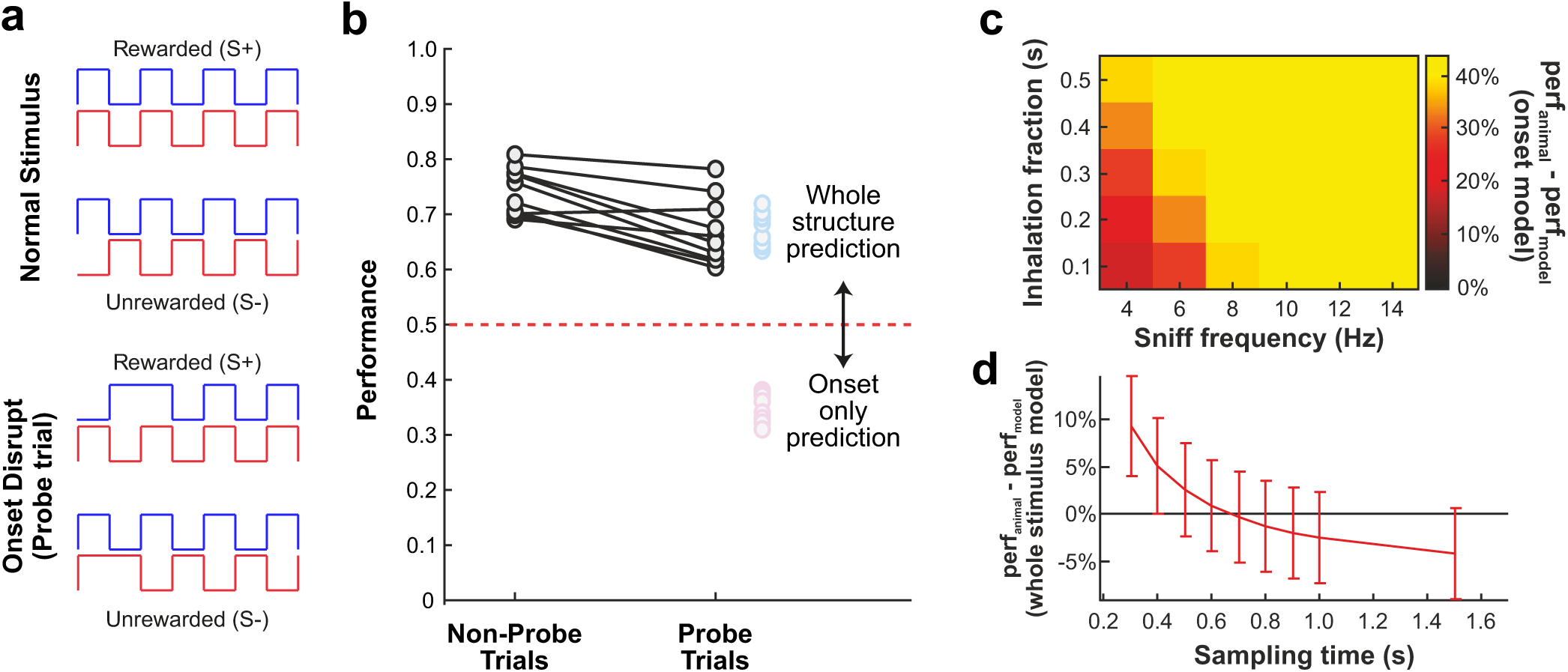
Onset detection. **a,** One explanation for the ability of mice to discriminate odour correlation structure could be that they sensitively detect simultaneity of odour onset with high temporal resolution rather than correlation structure across the stimulus. We therefore designed a new experimental stimulus in which the first stimulus pulse was disrupted to be presented on “catch trials”. Top: normal stimulus design, bottom: onset disrupt stimuli in which the first pulse in a correlated stimulus is disrupted to be anti-correlated; and vice versa for an anti-correlated stimulus. **b,** If animals solved the discrimination task by sampling the onset of the odour pulse only, their performance for onset disrupt stimuli should be ‘flipped’, i.e. should be significantly below chance since they would consistently make the wrong decision for both S+ and S-stimuli (pink dots). If they take the whole correlation structure into account, performance is predicted to be slightly degraded (as the beginning of the stimulus is “wrongly” correlated) but significantly above chance (blue dots). Animals were trained on standard (non-probe) correlation discrimination trials (f = 10 Hz) but onset disrupt (probe) trials were presented randomly on catch trials with a 1/10 probability. Performance was indeed only slightly degraded on probe trials but not flipped suggesting animals sampled multiple stimulus pulses to solve the discrimination. **c,** Heatmap of difference between onset model and actual animal performance across a range of sniff frequencies and inhalation fractions (n = 10). **d,** Difference between whole stimulus model and animal performance across different stimulus sampling times (n = 10, mean +/−SD).

**Supplementary Figure 2.5:**
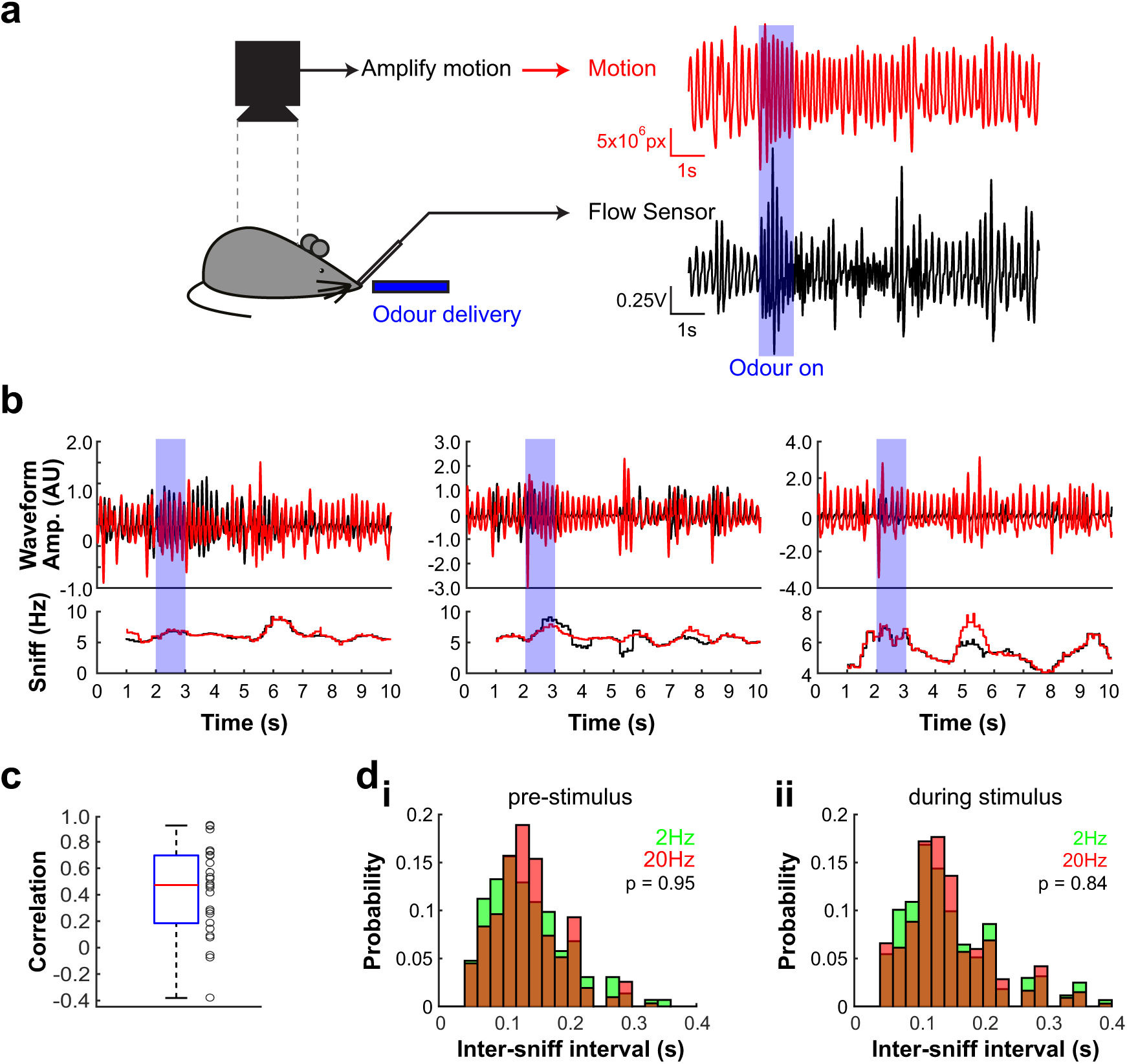
Respiration recordings. **a,** An overhead camera was used to image a head fixed mouse during a sequence of odour presentation. Simultaneously, a flow sensor was placed close to one nostril to monitor respiration to establish the validity of motion imaging-based respiration recording. Phase-based motion amplification was used to magnify motion on the animal’s flank to capture body movements associated with respiration. Right: example for simultaneous respiration measurement with motion imaging and flow sensor (see supplementary movies). **b,** Three further example trials with respiration rate extracted by motion imaging (red) and simultaneous flow sensor recording (black). Below: instantaneous sniff frequencies calculated from either sensor were tightly correlated. **c,** Correlation between respiration traces extracted motion imaging and respiration captured by flow sensor (n = 26 trials, 10 s duration each). Box plot shows median (red line), 25th to 75th percentile (blue box). **d,** Probability distributions of inter-sniff intervals for odour presentations (Amyl acetate vs Ethyl butyrate, 2 Hz and 20 Hz) for freely moving animals in AutonoMouse before stimulus onset **(di)** and during 2 s odour stimulation **(dii)** (n = 605 sniffs for 2 Hz and n = 668 for 20 Hz).

**Supplementary Figure 3.1:**
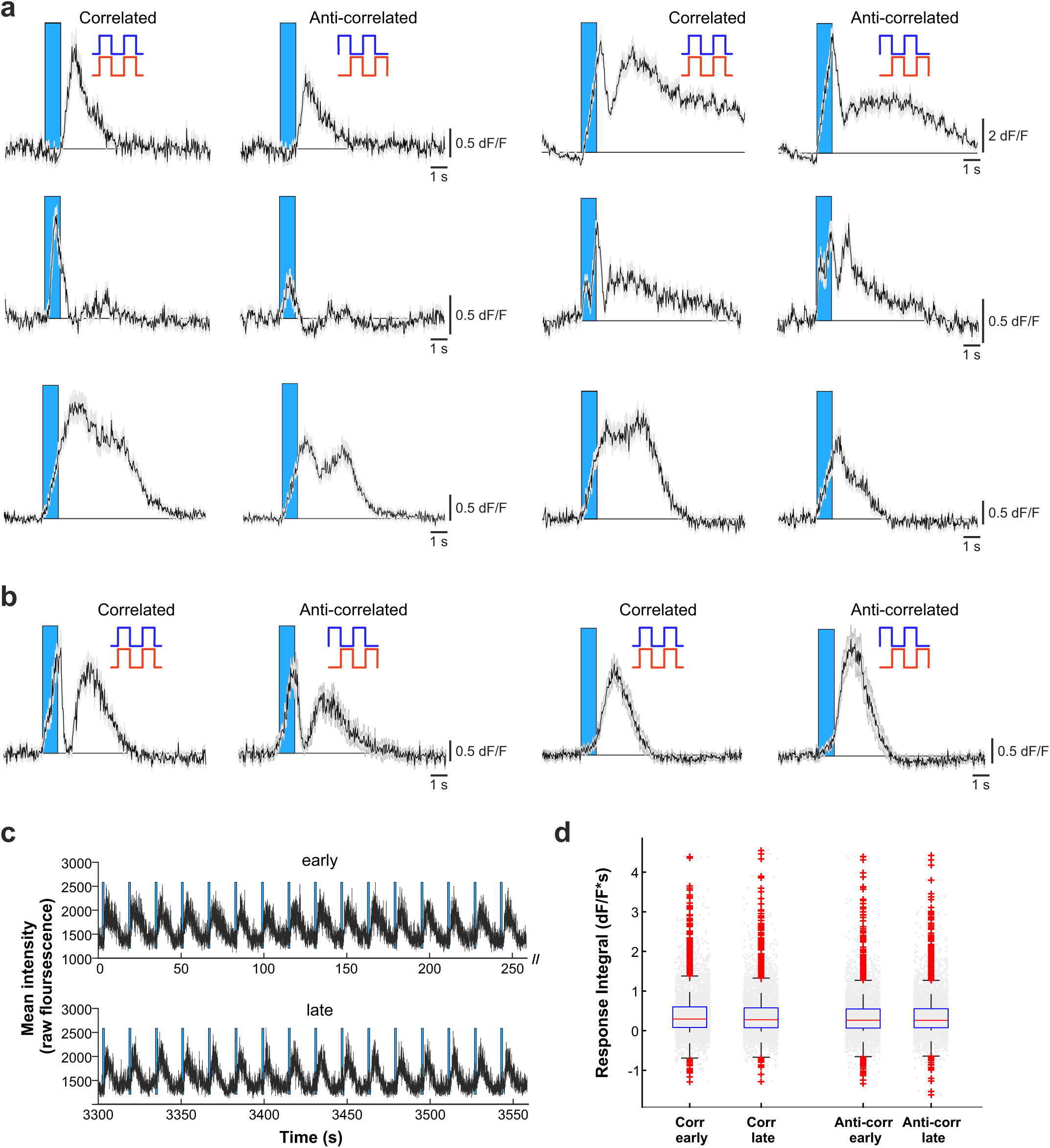
Additional example cells and signal stability. **a,** Example traces of ROIs that show differential response kinetics to correlated (left) and anti-correlated (right) stimulation (mean of 24 trials +/−SEM, f = 20 Hz). **b,** Example traces of ROIs that show differential response kinetics to correlated (left) and anti-correlated (right) stimulation (mean of 16 trials +/−SEM) in an awake head-fixed animal (mean of 24 trials +/−SEM, f = 20 Hz). **c,** Mean intensity of an example ROI over the course of an approximately 60 min imaging experiment. Shown are responses to the first (early) and the last (late) 16 odour stimulations, 1 s stimulation period is marked with blue bars. **d,** Summary of integrals of the 5 s response windows of all ROIs (n = 661) during early and late correlated and anti-correlated odour stimulations.

